# Dragonfly Inspired Smart Soft Robot

**DOI:** 10.1101/2020.04.28.067033

**Authors:** Vardhman Kumar, Ung Hyun Ko, Yilong Zhou, Jiaul Hoque, Gaurav Arya, Shyni Varghese

## Abstract

Recent advancements in soft robotics have led to the development of compliant robots that can exhibit complex motions driven by living cells(*1*, *2*), chemical reactions(*3*), or electronics(*4*). Further innovations are however needed to create the next generation of soft robots that can carry out advanced functions beyond locomotion. Here we describe *DraBot*—a dragonfly-inspired, entirely soft, multifunctional robot that combines long-term locomotion over water surface with sensing, responding, and adaptation capabilities. By integrating soft actuators, stimuli-responsive materials, and microarchitectural features, we created a circuitry of pneumatic and microfluidic logic that enabled the robot to undergo user- and environment-controlled (pH) locomotion, including navigating hazardous (acidic) conditions. DraBot was also engineered to sense additional environmental perturbations (temperature) and detect and clean up chemicals (oil). The design, fabrication, and integration strategies demonstrated here pave a way for developing futuristic soft robots that can acclimatize and adapt to harsh conditions while carrying out complex tasks such as exploration, environmental remediation, and health care in complex environments.

---

Soft bodied invertebrates and vertebrates have been an inspiration for the development of soft robots(*5*, *6*). Compared to robots made from hard components, soft robots have the ability to carry out sophisticated and delicate tasks inside confined spaces and complex environments(*7*–*9*). Such robots could be deployed into the real world to aid humans with tasks such as environmental monitoring, exploration, and remediation, or detection and handling of biological and chemical entities(*10*, *11*). In doing so, these robots must exhibit efficient locomotion, rapid responsiveness to environmental changes, and adaptation akin to living organisms while navigating natural environments with immense diversity. Most research in the area of soft robotics has focused on mimicking zoomorphic motions, including bending(*12*–*15*), crawling(*4*, *13*), climbing(*16*), rolling(*17*), jumping(*18*) and swimming(*1*, *2*, *14*, *19*), inspired by organisms such as snakes, octopuses, spiders, caterpillars, jellyfish, and stingrays. To enable these robots to perform more complex functions, such as sensing, reporting, and long-distance locomotion, researchers have had to incorporate either hard electronics(*20*), which compromises the compliance of the robots and their compatibility in aqueous or corrosive environments, or living entities such as living cells, which limits the life and resilience of the robots(*21*).

Here we describe a new class of completely soft and synthetic robots that display spatiotemporal control over travel distance, speed, and direction and can also respond and adapt to environmental changes and report encountered perturbations. In particular, we developed a dragonfly-inspired soft robot “DraBot” (**Fig. 1A**) that mimics the skimming ability of dragonflies on water surfaces over large distances. Long-distance locomotion in DraBot was enabled by externalizing the propellant reservoir and user control over the speed and direction of travel was executed by a cascade of pneumatic and microfluidic logic gates. Besides user control, DraBot was designed to exhibit pH-driven control over locomotion by utilizing spatially-confined pH-responsive features, which gave the robot additional environmental sensing and adaptation capabilities. Additional features, namely temperature sensing and oil detection, were achieved though the incorporation of thermochromic pigments and hydrophobic structures with microarchitectures.

**Fig 1.**
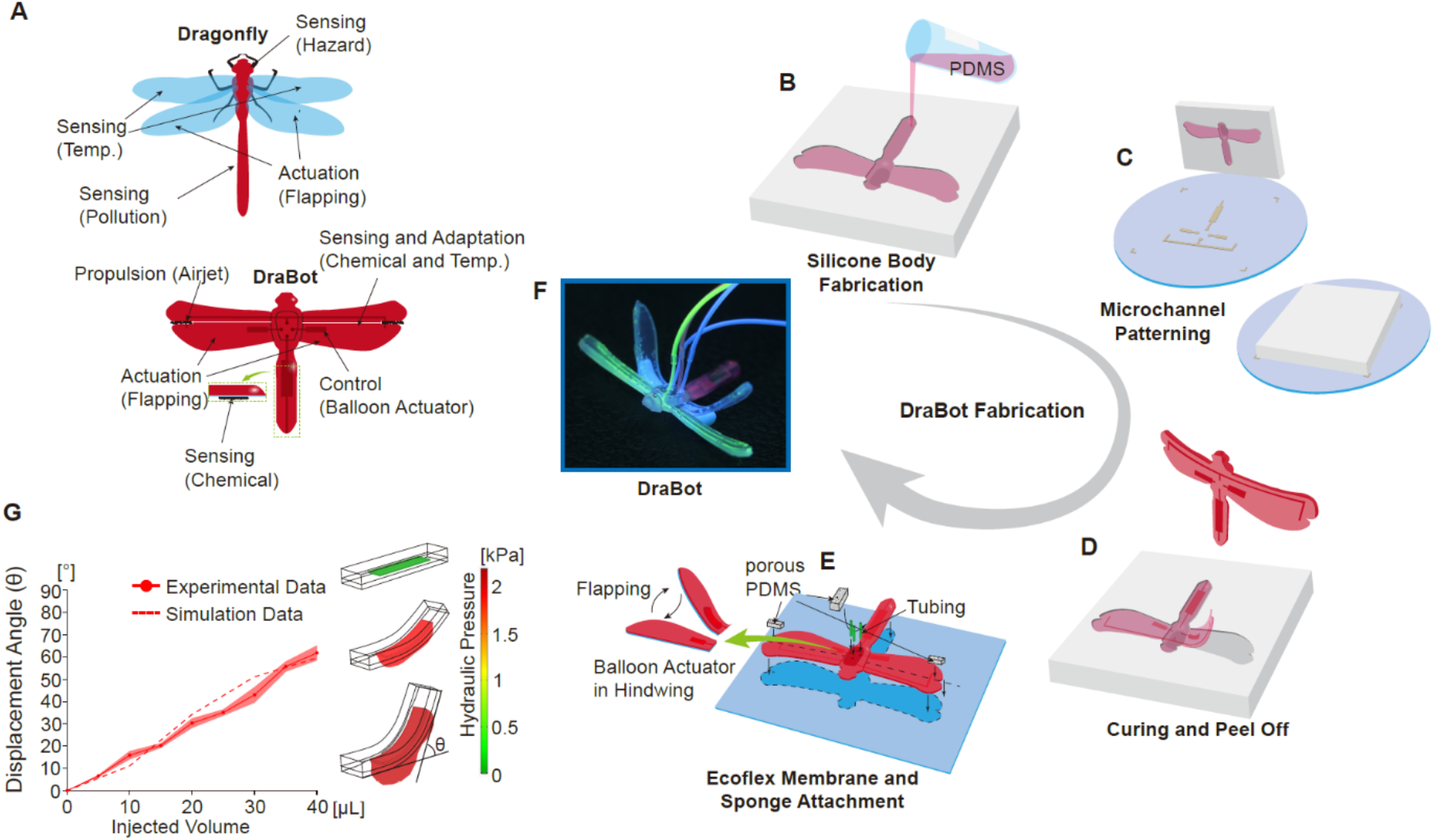
Design and fabrication of DraBot. (**A**)Structural and functional features of DraBot emulating a dragonfly. Schematic of the fabrication process: (**B**) pouring of PDMS precursor on the aluminum mold, (**C**) inversion onto the patterned silicon mold with microfluidic channels, (**D**) peeling off the DraBot body after PDMS curing, (**E**) and attachment of Ecoflex membrane, porous PDMS, and silicone tubings. (**F**) Digital image of DraBot with fluorescent ink in the microfluidic channels. (**G**) Comparison of theoretical predictions (dashed line) and experimental measurements (solid line) of balloon actuator-mediated deflection of the hindwing as a function of the volume of water injected into the hydraulic channel. The images on the right show the development of pressure inside (increasing from top to bottom) the balloon leading to upward flexing of the wing.

The base of DraBot was fabricated from silicone elastomers with a multi-step fabrication process as detailed in Supplementary Informations(**Fig. 1B-F**). Briefly, poly(dimethyl siloxane) (PDMS) was cured to achieve dragon fly-specific structural features embedded with microfluidic channels (**Fig. 1B-D**) and capped with Ecoflex membrane (**Fig. 1E**). The forewings were embedded with pneumatic channels to drive propulsion, and the hindwings with hydraulic channels leading to microfluidic balloon actuators, which regulate the “flapping” (flexing) angle at the base of the hindwings for controlling locomotion. The optimal channel dimensions for achieving the targeted flapping behavior were identified from a combined finite element analysis of fluid dynamics and bending mechanics using experimentally determined material properties as input parameters (**Fig 1G**, **Fig. S1** **and** **S2**, **and Movies S1 and S2)**. The hindwings and the abdomen of DraBot were decorated with PDMS structures containing interconnected micropores (**Fig. 1E** **and** **Fig. S5**). These porous structures offer multiple functions: flotation and motion control in combination with the balloon actuators. Finally, air and fluid soft tubings were connected to the corresponding pneumatic and hydraulic channel inlets for user-controlled locomotion (**Fig. 1E**) to yield the final functioning robot (**Fig. 1F**).

DraBot locomotion is driven by pneumatic propulsion through microchannels in the forewings by supplying compressed air via external tubing. The outlets of the pneumatic channels in the forewings face the hindwings allowing the pressurized air to exit in the backward direction, enabling the motion to be controlled by the positioning of the hindwings. In the rest state, the hindwings are coplanar to the forewings, causing the porous PDMS structures on the left and right hindwings to block the air outlets on the forewings. This causes the exiting airflow from the forewings to be dispersed in random directions through the micropores of the PDMS substrate, resulting in no net propulsion (**Fig. 2A,C**). To enable forward propulsion, both hindwings were bent upwards, through the action of balloon actuators, which moves the blocking PDMS structure out of the way of the air outlets in forewings (**Movie S3**). The unidirectional air flow now provides a net forward propulsion to the bot, making it move in the forward direction (**Fig. 2B,D**). The speed of the robot could then be simply controlled by the input pressure into the pneumatic channel; **Fig. S3** shows the velocity of the DraBot as a function of input air pressure. For all locomotion studies presented here, we employed 30 psi as the input pressure, resulting in a velocity of ~10 cm/s.

**Fig. 2.**
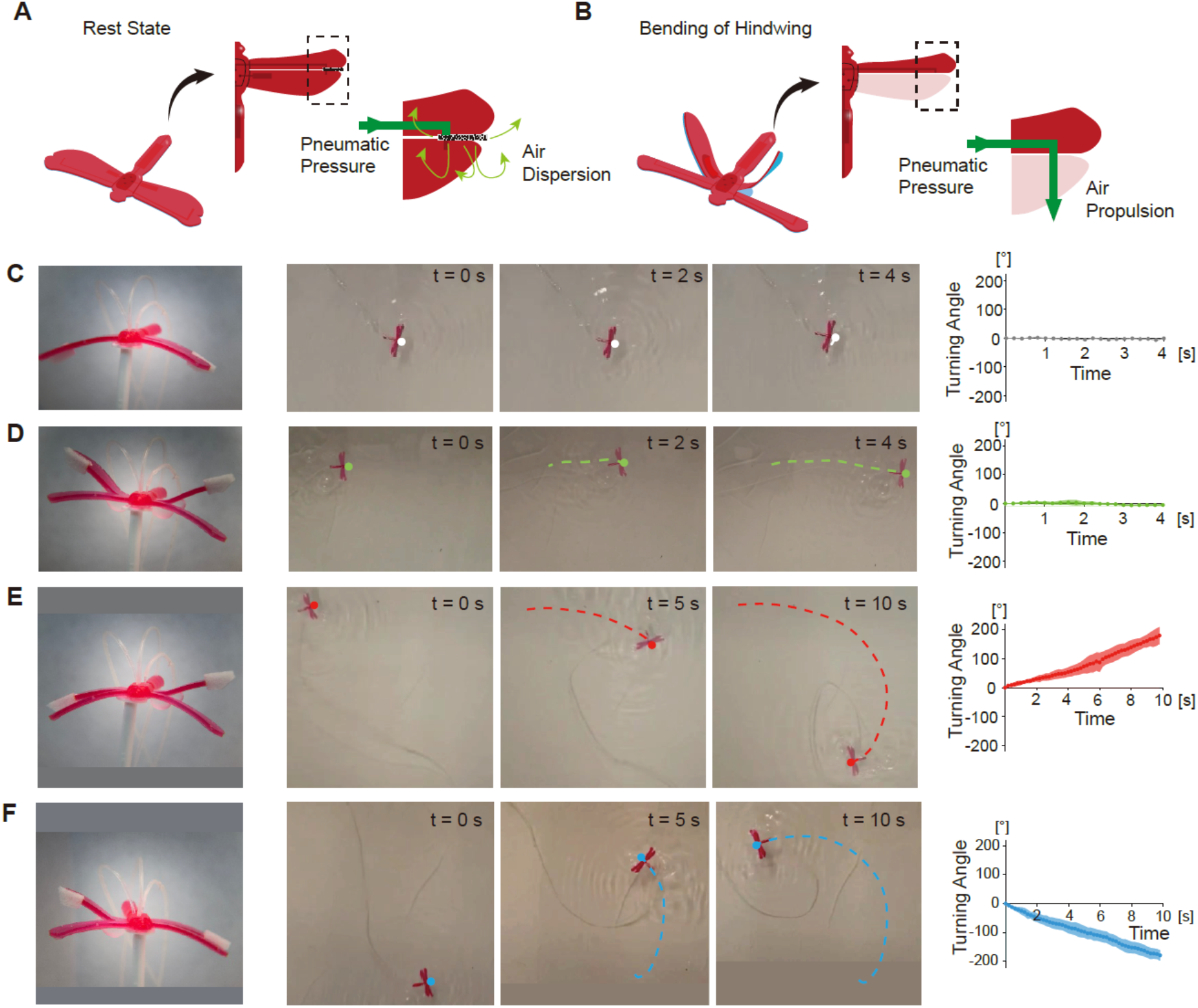
User-controlled locomotion and maneuvering. Schematic depicting the conformation of the hindwings and corresponding air flow at the pneumatic channel exit in case of (**A**) resting and (**B**) upward-flexed hindwings. (**C**) Resting state of both hindwings results in no net movement of DraBot. (**D**) Upward flexing of both hindwings leads to straight motion. (**E**) Upward flexing of left wing results in a right turn, and maintaining this flexed state results in clockwise motion. (**F**) Upward flexing of right wing results in a left turn, and maintaining this flexed state results in counter-clockwise motion.

The bot maintains a straight path over time without any turns when undergoing forward motion (**Fig. 2D** **and Movie S4**). To introduce changes in the direction of motion, we used the balloon actuators to flap the hindwings, which misaligns the microporous PDMS substrate and the air outlets. Specifically, to make the bot turn right, the left hindwing was flapped upwards, which exposed the left air outlet, generating a net torque resulting the bot to turn rightwards (**Fig. 2E**), and *vice versa*, flapping the right hindwing caused the bot to turn leftwards (**Fig. 2F**). The extent to which the bot changes direction depends on the duration of time a wing is flapped upwards. A sustained left- or right-turn signal led to continuous circular motion (**Movies S5 and S6**). These precise controls over locomotion and direction allows the robot to be used to explore water surfaces (**Movie S7**).

The entire control system of DraBot can be mapped onto a cascade of logic gates that includes 4 AND, 2 XOR, and 1 OR gates (**Fig. 3A**). The user input is comprised of three Boolean signals – one pneumatic signal (*P*), that diverges to both forewings, and two hydraulic signals (*H*_L_ and *H*_R_) for the balloon actuator in the left and right hindwings – which we denote using [*P*, *H*_L_, *H*_R_]. There are three outputs, one corresponding to forward motion (*F*) and the other two corresponding to left (*T*_L_) or right turn (*T*_R_), denoted by [*F*, *T*_L_, *T*_R_]. The resting position of hindwings corresponds to [0,0,0] or [1,0,0] inputs and leads to an output of [0,0,0]. The straight motion of the bot corresponding to [1,0,0] is achieved with an input of [1,1,1] corresponding to both wings flapped up. The left and right turning motions corresponding to outputs [0,1,0] and [0,0,1] are achieved through [1,0,1] or [1,1,0] inputs, respectively (**Fig. 3B**).

**Fig. 3.**
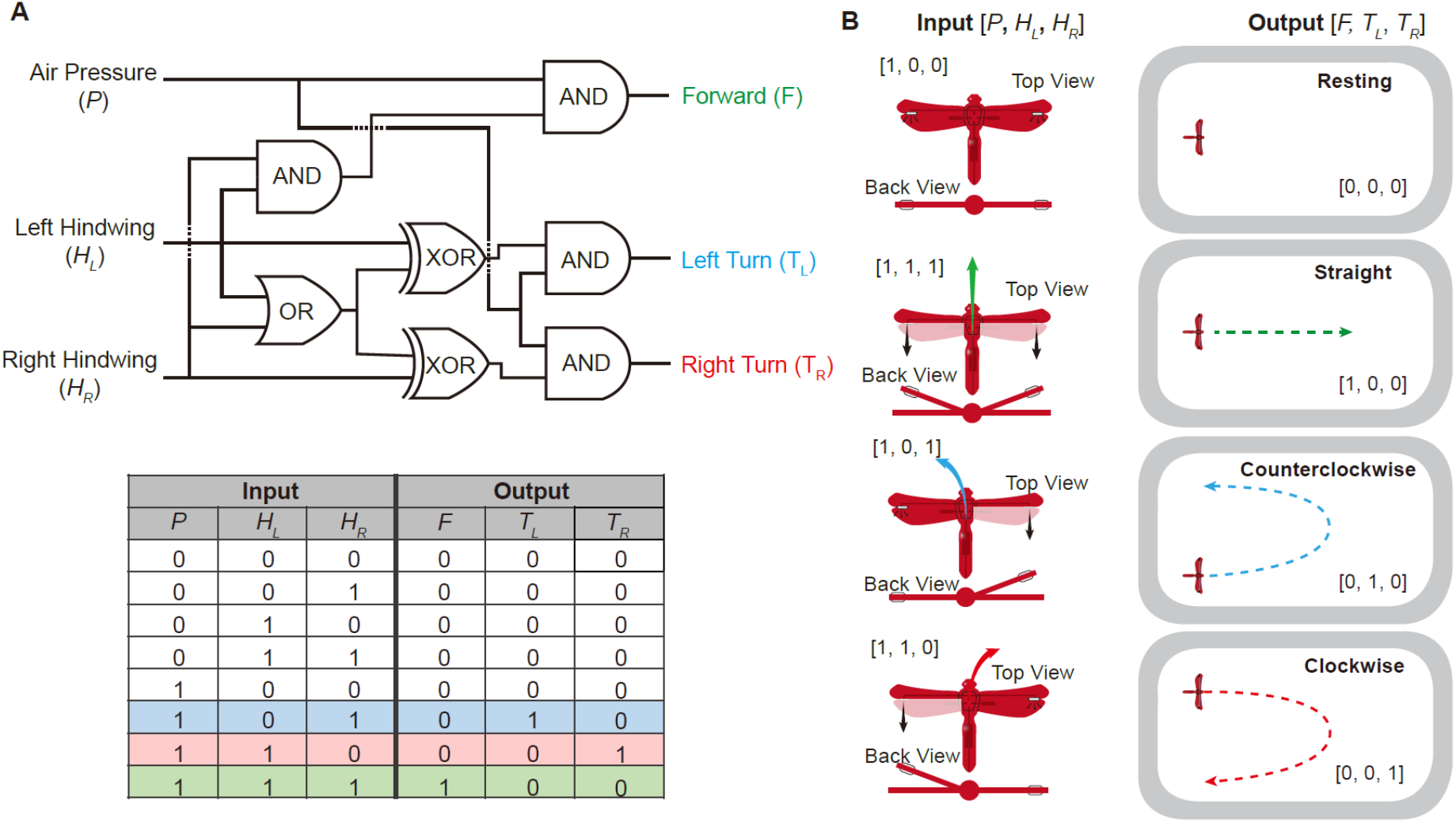
Logic diagram of user-controlled locomotion. (**A**) Logic diagram and truth table describing user control over DraBot locomotion and maneuvering. (**B**) Schematic mapping of the input signals to the corresponding output motion of the DraBot.

While the above design principles can introduce long-term locomotion into a soft robot, the motion is still entirely user controlled. However, living organisms not only exhibit locomotion, but can also sense their environment and adapt their motion accordingly while simultaneously carrying out other functions. We further engineered DraBot to simultaneously exhibit several of these characteristics of living systems. First, we introduced environment-adaptive motion by designing the wings on one side to be responsive to pH changes. To this end, a surface cut was introduced across the right fore- and hindwings, and the “wound” was “dressed” with a acryloyl-6-aminocaproic acid (A6ACA) hydrogel that shows instantaneous reversible pH-responsive healing(*22*) (**Fig. 4A**, **Fig. S4**). At low pH, the A6ACA hydrogel welds the two left wings together, preventing the hindwing from flapping, and thereby blocking the air exit channel on the left forewing (**Fig. 4B**, **Movie S8**). Thus, when the bot encounters acidic conditions, it makes a continuous left turn (resulting in counterclockwise circular motion), despite receiving a user input signal for forward motion (**Fig. 4C**, **Movie S9**). Restoring neutral conditions dissociates the welded state of the right wings and the bot resumes its original forward motion that it exhibited before encountering the acidic conditions. Hence, through judicious incorporation of stimuli-responsive materials into DraBot, it was able to not only sense and adapt to environmental pH changes in water, but also report on such perturbations via observable changes in motion. Second, we mimicked the thermo-responsive color changes in dragonflies(*23*, *24*) by encoding DraBot’s wings with a thermochromic pigment which changes color in response to temperature changes, wherein the wing color changed smoothly from red to yellow with increase in temperature (**Fig. 4D**, and **Movie S10**). Lastly, the hydrophobicity along with the large surface area of the microporous PDMS structure at the abdomen and wings of DraBot was further exploited to detect and absorb hydrophobic entities such as oils from the surface of water (**Fig. 4E** and **Movies S11 and S12**). Efficient oil absorption is achieved through the coupled action of hydrophobicity and capillary forces resulting from the 3D microporous structure of PDMS.

**Fig. 4.**
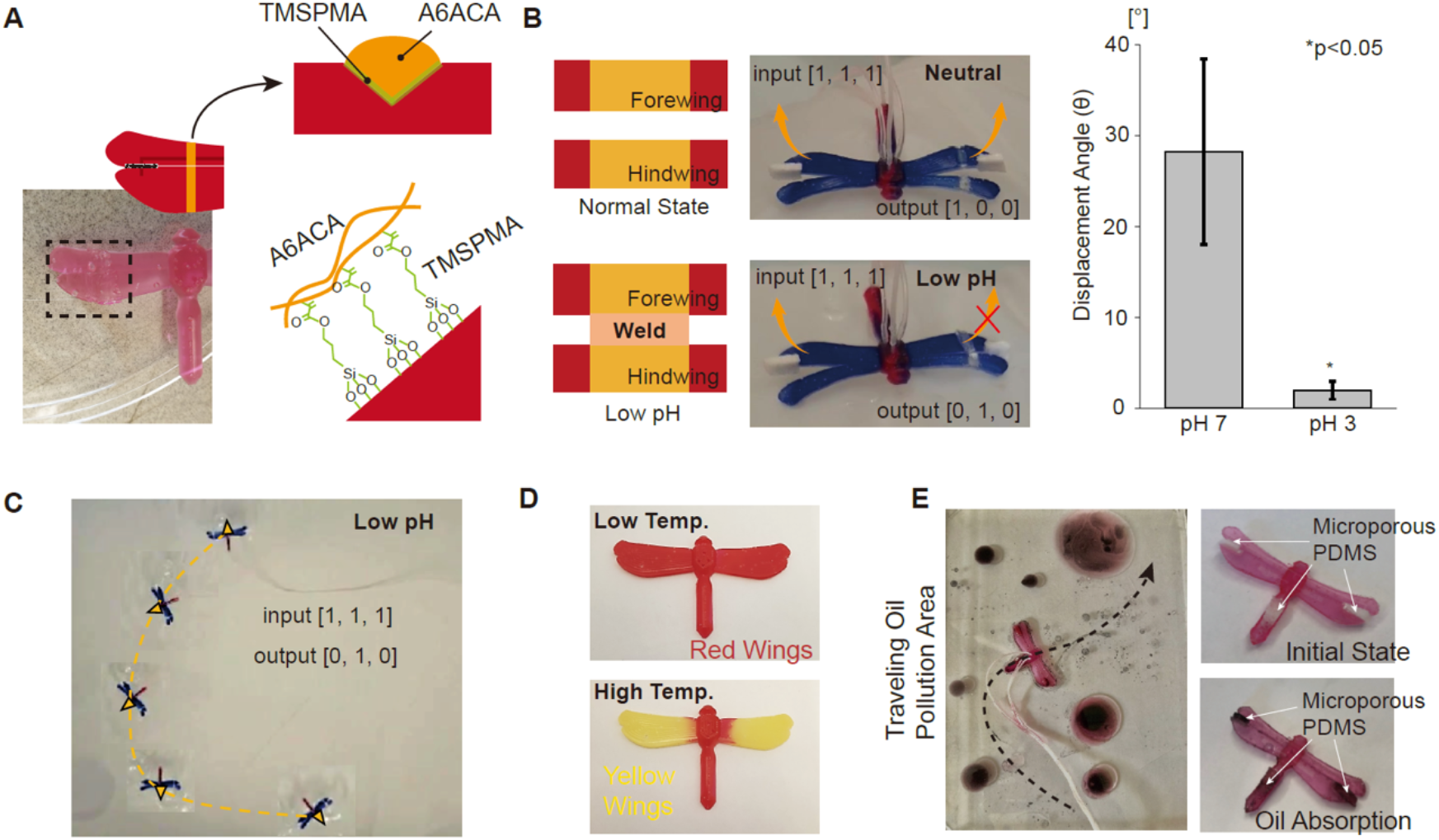
Multifunctionality of DraBot. (**A**) Modification of the left wing to achieve pH sensing and environment-driven motion. (**B**) pH-mediated healing of A6ACA hydrogel coating on left fore- and hindwing in acidic condition welds them together and disables flapping. The left column provides a schematic depiction of un-welded and welded left wings at normal and low pH environments, respectively. The top and bottom images in the middle column show the digital images of DraBot at corresponding pH states. The right column provides the corresponding flexing angle of the left wing in two states. (**C**) Counter-clockwise motion of DraBot over time in acidic condition depicted by overlaying the images of the bot at different time points. (**D**) Changes in wing color at different temperatures. The wings change from red color at room temperature (top image) to yellow above 37 °C (bottom image). (**E**) Color change of domains containing porous PDMS substrates after traveling through water polluted with colored oil. The left image shows DraBot traveling through the water contaminated with oil. The right images show the domains before (top) and after (bottom) encountering oil contaminants.

In summary, we have described the development of a completely soft, multifunctional robot that travels large distances across the surface of water in a user- and environmentally-controlled manner. This was achieved by integrating hydraulic-pneumatic logic with environment-responsive smart materials. A key design strategy to achieving such multifunctionality is the incorporation of multiple detection mechanisms, each confined within distinct structural segments of the robot, which enables simultaneous detection and reporting of multiple environmental stimuli or hazards. For instance, the ability of the robot to respond to pH can be leveraged to detect freshwater acidification, which is a serious environmental problem affecting several geologically-sensitive regions(*25*, *26*). Similarly, the temperature-sensitive color change of the robot’s wings can be used to identify changes in the surface temperature of water associated with red tide(*27*), bleaching of coral reefs(*28*), and decline in population of marine life(*29*). Local increase in surface temperature and pH drop of ocean water are also indicators for submarine volcanic activities(*30*) that can trigger earthquakes and tsunamis. Finally, the ability of the microporous PDMS substrates to absorb hydrophobic moieties makes such long-distance skimming robots an ideal candidate for early detection of oil spills. Such spills are indeed a serious environmental and economic hazard as evident from the deep horizon oil spill in 2010(*31*, *32*). Besides detection, one could also envision using smart robots to clean-up oil spills or stop leakages by sealing cracks responsible for the spill. In addition to exploring the environment, smart soft robots could also find applications in healthcare(*33*) where the robots could be used to diagnose contagious diseases and to aid frontline health care workers in providing care to patients(*34*).

## Acknowledgements

This work was performed in part at the Duke University Shared Materials Instrumentation Facility (SMIF), a member of the North Carolina Research Triangle Nanotechnology Network (RTNN), which is supported by the National Science Foundation (Grant ECCS-1542015) as part of the National Nanotechnology Coordinated Infrastructure (NNCI);

## Author contributions

V.K. and S.V. conceived and designed the overall research. U.H.K. and V.K. performed experiments and data analysis. Y.Z. and G.A. performed all the simulations and associated analysis. J.H. carried out all chemical synthesis and SEM imaging. V.K., U.H.K., G.A., and S.V. wrote the manuscript with contributions from all authors;

## Data and materials availability

All data is available in the main text or the supplementary materials.

## Supplementary Materials

### Materials and Methods

#### Finite element analyses to determine hydraulic channel dimensions for balloon actuators

A 3D numerical simulation was developed in COMSOL with an input of experimentally determined material properties to design the balloon actuators, which were used to control the bending angles of DraBot’s hindwings. The inflation of the balloon actuator induces a pulling force on the microfluidic channel, which when strong enough can bend the channel structure and thus generate an out-of-plane motion(*35*). The extent of bending achieved highly depends on the material properties of the two material layers sandwiching the channel, as also reported by previous works(*35*, *36*). The technical details of the model has been described in the Supplementary Text.

#### Mechanical properties of Ecoflex

Hyperelastic constants of the Ecoflex 00-30 were determined by tensile measurements using an ElectroForce 3220 Series III (TA Instruments, Inc.). Type V (i.e, dumbbell) samples with 4 mm thickness were prepared according to ASTM (American Society for Testing Materials) standard (**Fig. S1A**). The stress-strain measurements were carried out with a load cells of 225 N at 0.25 mm/min. The two-term Ogden model described below was used to fit the measured stress strain curve to obtain the hyperelastic constants (**Fig. S1B** **and** **S1C**) employed in our balloon actuator simulations.

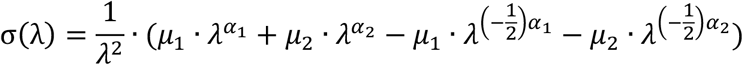

where,

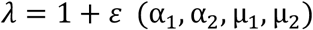

where *λ* is the principle stretch and α_1_, α_2_, μ_1_ and μ_2_ are material constants.

#### Experimental validation of theoretical predictions

To validate the theoretical prediction from COMSOL simulations, rectangular microfluidic channels were prepared with constant channel width × height (1.8 mm × 0.15 mm) and varying length (4.1, 5.6, and 7.1 mm) using soft lithography. The patterned silicon wafer was made using SU-8 photolithography. Sylgard 184 PDMS was poured onto the patterned silicon surface and cured at 60 °C for at least 2 hours. Cured PDMS was released from the wafer, and a 1 mm inlet was created using a biopsy punch (Miltex, Inc.). Each actuator was pressurized by a syringe pump (Harvard Apparatus, Inc.) at a flow rate of 2.5 μl/s, and the angular displacement corresponding to the injected fluid volume was measured. Three cycles for each actuator were tested for data acquisition. Channels with a dimension of 7.1 × 1.8 × 0.15 mm (length × width × height) were used for all further studies.

#### Soft Robot Fabrication and Assembly

The aluminum mold with dragonfly structural features (3.5 cm wingspan and 5.7 cm length) was fabricated by CNC machining (Xometry, Inc.) with aluminum 6061-T6 material. Pneumatic and hydraulic channels were fabricated using soft lithography wherein patterns with a height of 150 μm were etched using an SU-8-100 photoresist (MicroChem, Inc.) on a silicon wafer (100 mm, (1 0 0), boron-doped, p-type, ID: 452, University Wafer). Detailed experimental steps for patterning of silicon wafer through baking, aligning, exposing, and developing were followed as per the specifications in the product datasheet. This patterned silicone wafer was used as a master mold for the pneumatic and hydrolytic channels. Sylgard 184 (PDMS; Dow Corning, Corp.) was mixed at 10:1 ratio (base : curing agent), degassed, and poured onto the silicon wafer and aluminum mold. The mold was the inverted onto the silicon wafer to encase the PDMS between the mold and the silicon wafer and cured at 60 °C for two hours. The cured Drabot was peeled off from the mold, 1 mm diameter channel inlets (Miltex, Inc.) were punched, and the bot was attached on 0.5 mm thick Ecoflex 00-30 silicone rubber membrane (Smooth-on, Inc.) using O_2_ plasma (K1050X, Quorum Technologies, Ltd.) at 50 W for 30 s. The membrane was attached to the body immediately after the plasma treatment and placed on a hotplate at 200 °C for 5 min followed by baking at 60 °C for two hours to ensure strong and durable bonding. Flexible silicone tubing, with an internal diameter of 0.0635 cm (McMaster-CARR, Inc.), was inserted into the chest area of the dragonfly-bot at inlets of the pneumatic and the two hydraulic channels.

#### PDMS substrate with interconnected microporous structure

Microporous PDMS substrates were prepared using salt leaching method. Granular salt (NaCl) was filled in a 24 well plate and exposed to a humid environment. Upon hardening and clump formation, 10:1 PDMS mixture (base: curing agent) was poured onto wells containing NaCl clumps, and was placed in a vacuum desiccator for 4 hours to allow the PDMS solution to penetrate into the salt clumps. The PDMS infiltrated clumps were allowed to cure at 60 °C for 2 hours and then immersed in water overnight on an orbital shaker to dissolve the salt. The dissolution of the solid salt resulted in a PDMS with interconnected pores(*37*, *38*). The morphology of the PDMS substrate formed was analyzed by SEM. A PDMS slab and microporous PDMS were cut into smaller pieces (~3 cm × 3 cm × 3 cm), and sputter coated with gold for about 90 sec at 12 mA current using Denton Vacuum Desk V Standard sputter coater. The gold coated samples were imaged using FEI Apreo scanning electron microscope at 2 kV operating voltage. Images were acquired from at least three different positions (**Fig. S5**). The microporous PDMS substrate was cut into desired shapes and attached at hindwings and abdomen site of the DraBot.

#### Synthesis of *N*-acryloyl 6-aminocaproic acid (A6ACA)

*N*-acryloyl 6-aminocaproic acid (A6ACA) was synthesized from 6-aminocaproic acid (6ACA) (Sigma Aldrich, Inc.) as described elsewhere(*39*). Briefly, 0.1 M 6ACA and 0.11 M NaOH were dissolved in 80 mL deionized water in an ice bath with constant stirring. Following complete dissolution, 0.11 M acryloyl chloride (Sigma Aldrich, Inc.) in 15 mL tetrahydrofuran was added dropwise. The pH was maintained at 7.5-7.8 by the addition of 2.5 M NaOH solution until the completion of the reaction. The reaction mixture was then acidified to pH 3 using 6 M HCl and extracted with ethyl acetate. The ethyl acetate layer was collected and dried over anhydrous sodium sulfate to remove any traces of water. After filtration of sodium sulfate, the solution was concentrated using a rotavapor and further precipitated in hexane. Precipitation was carried out in an ice bath under constant stirring. Repeated precipitation was used to purify the product, and the product was dried in a desiccator.

#### Surface Functionalization of PDMS

For the bonding of A6ACA hydrogel to PDMS, the PDMS surface was functionalized with 3- (Trimethoxysilyl) propyl methacrylate (TMSPMA) (Sigma-Aldrich Inc.). The PDMS slabs (wing of the DraBot) were cleaned by using sonication with isopropanol, ethanol, and deionized water for 5 min, respectively, and dried before use. The cleaned PDMS was subjected to oxygen plasma treatment (100W, K1050X, Quorum Technologies, Ltd.) for 3 mins, following which the PDMS was dipped into 2 wt % of TMSPMA in acetic acid solution (pH 3.5, dissolved in deionized water) for 2 hours at room temperature. The PDMS was then washed with ethanol, dried, and stored in low humidity conditions until used.

The functionalization was characterized by Fourier Transform Infrared Spectroscopy (FT-IR). FTIR analyses was performed on 1 cm x 1 cm PDMS substrates with a thickness of 1 mm to examine the effect of oxygen plasma and TMSPMA modification. The infrared spectrum was detected using Thermo Electron Nicolet 8700 equipped with the ATR module. Each sample was scanned 32 times at room temperature and atmospheric pressure. The data (**Fig. S4**) were collected in the range of 4000 ~ 500 cm^−1^ and analyzed using OMNIC software (Thermo Fisher Scientific Inc.).

#### Coating of A6ACA hydrogel onto functionalized PDMS

Functionalized PDMS were coated with a thin layer of A6ACA precursor solution and polymerized via free-radical polymerization at 37 °C for 3 hrs. The precursor solution was prepared by mixing 0.1% *N*, *N′*-methylene bisacrylamide (Bis-Am) (Sigma-Aldrich, Inc.) with 1 M of A6ACA monomer dissolved in 1 N NaOH. Roughly, 0.5% ammonium persulfate (APS) and 0.15% tetramethylethylenediamine (TEMED) were used as initiator and accelerator, respectively.

#### Thermochromic Ink

Thermochromic pigment (Blue-Colorless or Red-Yellow, Atlanta Chemical, Inc.) was used for temperature sensing of DraBot. During the fabrication process, the pigment was mixed with PDMS before pouring into the wing region while the rest of the bot was fabricated out of PDMS mixed with regular silicone colorant (red).

#### Controlling the DraBot motion

The rear ends of the hydraulic tubings were connected to two 10 ml syringes controlled manually while the end of the air tubing was connected to the central compressed air. Thus, the span area of DraBot is *πl*^2^, for a tubing length of *l*. The exploring motions corresponding to various input pressures were recorded using a video camera and analyzed by ImageJ software (NIH). 30 psi air pressure was used as a standard input pressure for all experiments described in the manuscript.

### Supplementary Text

Details of the finite element model for balloon actuator is described below. Since the balloon is activated by water and the materials used for the channel are hyper-elastic, the simulation model considered complete coupling between fluid flow and solid deformation.

#### Governing equations

Fluid flow: As the gravitational force is negligible compared to the pressure generated by the water balloon, the time-dependent flow field in the microchannel is governed by the conventional continuity and momentum equations,

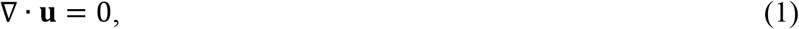

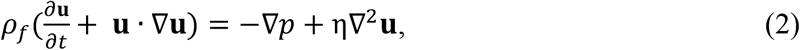

where *ρ*_*f*_, *p*, *η* and **u** are the fluid density, pressure, viscosity and velocity vector, respectively.

Solid mechanics: Since PDMS and the Ecoflex membrane endure large deformations, the second Piola-Kirchhoff stress tensor was used in the governing equations as,

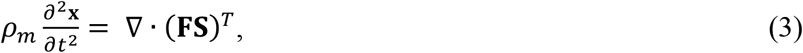

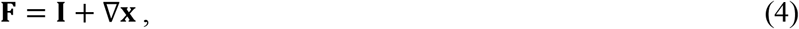

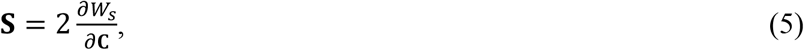

where **x** is the displacement vector, *ρ*_*m*_ is the material density, **F** is the deformation gradient, **S** is the second Piola-Kirchhoff stress tensor, **I** is the identity tensor, *W*_*S*_ is the strain-energy function, and **C** = **F**^*T*^**F** is the right Cauchy-Green deformation tensor. The Cauchy stress **σ** can be reconstructed in terms of **S**.

Since PDMS and Ecoflex membranes are hyperelastic materials, suitable strain-energy functions are needed to describe the nonlinear relationship between the stress and strain. For PDMS, the strain-energy function can be expressed by the Mooney-Rivlin model(*40*, *41*) as,

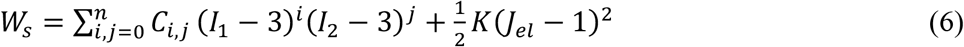

where *I*_1_ and *I*_2_ are the first and second invariant of the left isochoric Cauchy-Green deformation tensor, *K* is the bulk modulus, *J*_*el*_ is the elastic Jacobian, and *C*_*i,j*_ are material parameters.

In addition, the strain-energy function for Ecoflex membrane was represented by the Ogden material model(*42*) as,

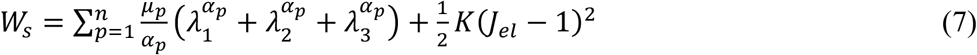

where *λ*_*i*_ (*i* = 1,2,3) are the principal stretches, and *μ*_*p*_ and *α*_*p*_ are material constants.

#### Computational domain and boundary conditions

The out-of-plane bending motion of the microchannel takes place in the lateral direction. Therefore, a 3D cuboid domain was chosen for the study set up as shown in **Fig. S2A**, where only a length of 12 mm in the lateral direction and 3 mm in the side direction were considered to capture the main dynamic motions of the channel so as to reduce computational cost. This model consists of two layers representing the PDMS and Ecoflex membrane. The fluid domain was embedded into the space between the PDMS and Ecoflex membrane layers, as shown in **Fig. S2A**, where the length *L* is an adjustable parameter. Note that the balloon expansion mainly appears in the rectangular fluid domain and the inlet tube used in the experimental device only serves as a passage to inject water into the domain; therefore, the part of tube has not been included in the computational domain. Thus, the fluid inlet has been directly assigned to the right end of the rectangular domain. The following boundary conditions were prescribed:

<Fluid flow> Inlet: volume flow rate *V*_0_; All other walls: no slip with **u** · **n** = 0; Initial conditions: **u** = 0 and *p* = 0.
<Solid mechanics (PDMS and membrane)> Right ends: fixed constraint with **x** = 0; All outer boundaries: free boundaries with zero stress. Initial conditions: **x** = 0 and 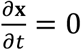.

#### Multiphysics

This fluid-structure interaction was assigned to the interfaces between the fluid domain and solid mechanics.

#### Numerical approach

The dynamic bending motion of the cuboid microchannel in Fig. 1 was simulated in COMSOL 5.3a. Eqs. (1-2) were solved in the Laminar Flow module, and the Moving Mesh was applied for the fluid domain. The Solid Mechanics module was used to solve for the structure deformation from Eqs. (3-7). Within the module, the five parameters Mooney-Rivlin material model (Eq.(6)) was used to calculate the strain-energy function for PDMS while Ogden material model (Eq. (7)) was used for Ecoflex membrane. The Mooney-Rivlin parameters for PDMS was used from a previous work(*43*). The Ogden material constants for Ecoflex00-30 membrane determined from the mechanical measurements were used in the simulation. The Fluid-Structure Interaction was assigned to the coupled interfaces between fluid flow and solid mechanics. A time-dependent study was added to solve the model. All the values of the physical and geometrical parameters are summarized in **Table S1**.

---

**Fig. S1.**
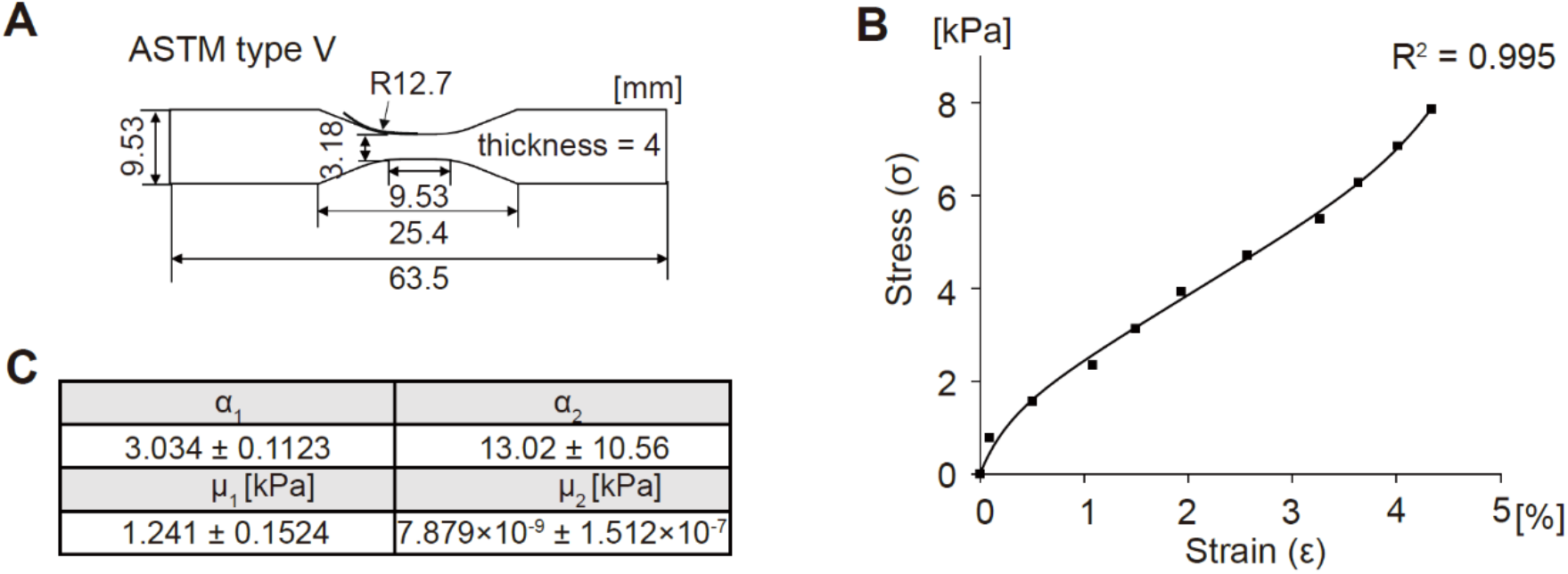
Mechanical test/analysis of Ecoflex 00-30. (**A**) Tensile test sample dimension according to the ASTM type V standard. (**B**) Curve fitting graph of Ecoflex 00-30 stress-strain data with Ogden model. (**C**) Obtained Ogden model parameters from curve fitting data.

**Fig. S2.**
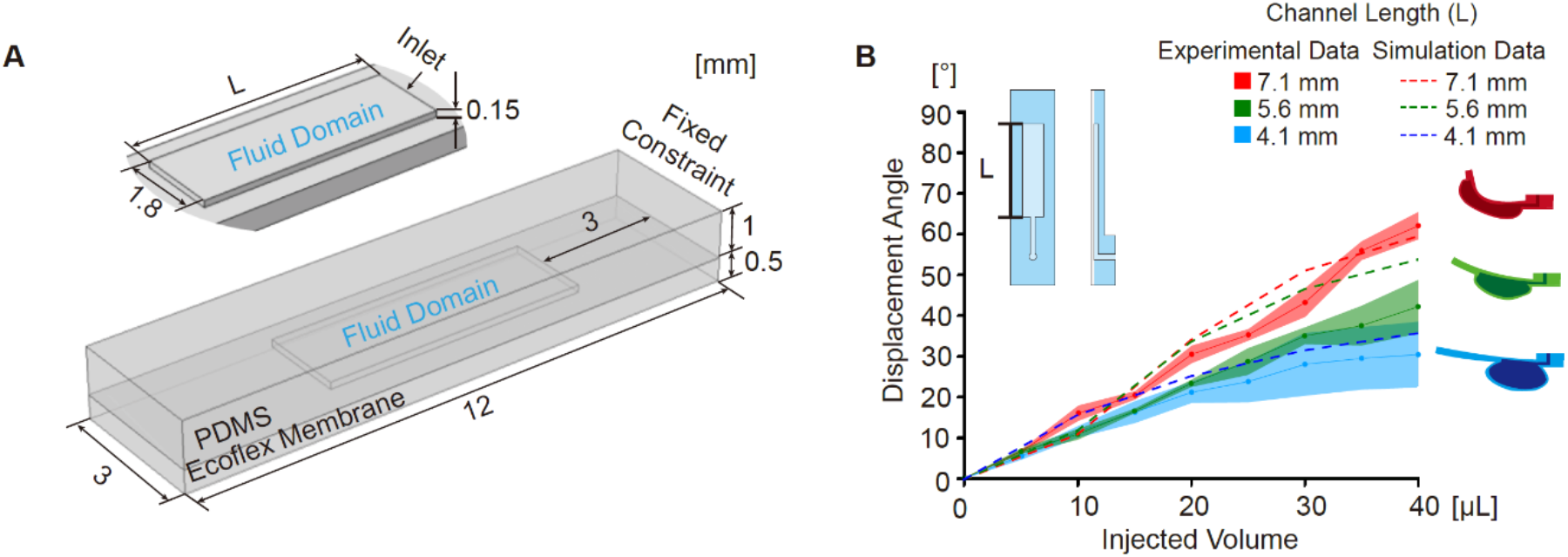
Simulation of balloon actuator for optimizing channel dimensions. (**A**) Isometric view of the 3D cuboid domain of balloon actuator. The fluid domain is embedded into the space between the PDMS and membrane layers, as highlighted in the inset. (**B**) Comparison of experimental (solid line) and simulation (dash line) results of displacement angle (θ) with different rectangular channel lengths: 4.1, 5.6, and 7.1 mm.

**Fig. S3.**
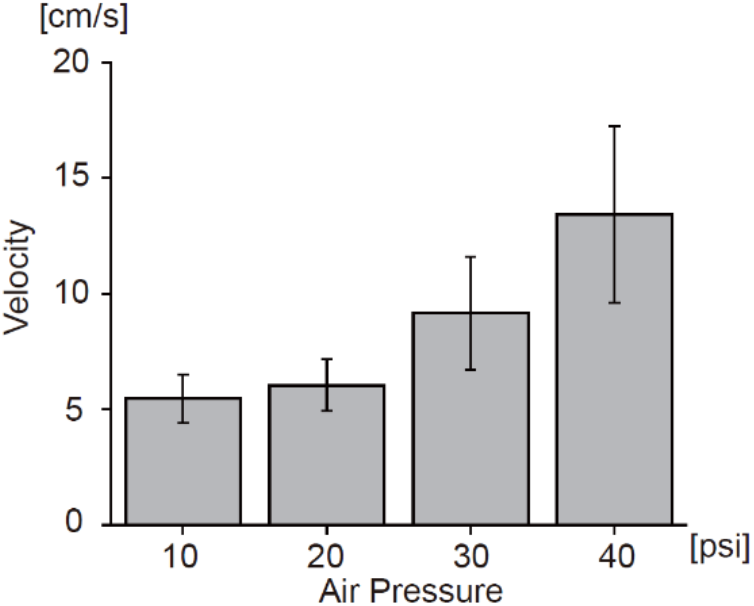
DraBot velocity in various pneumatic input pressure. Velocity increases from ~5.5 cm/s to ~ 13.4 cm/s when air pressure is increased from 10 to 40 psi.

**Fig. S4.**
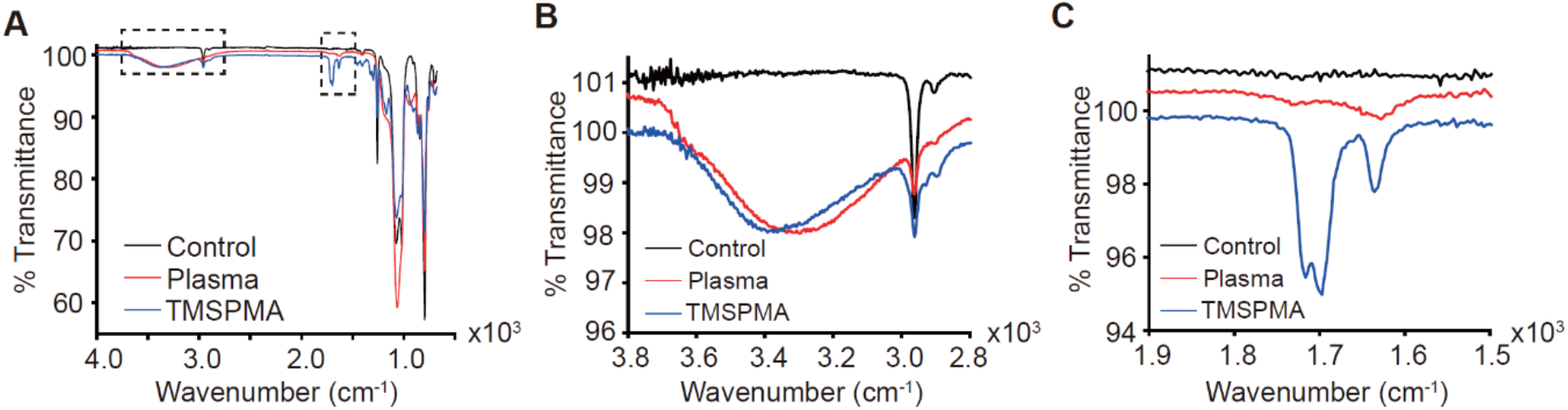
FTIR spectra of bare, plasma treated, TMSPMA modified PDMS surface. (**A**)The total FTIR graph in 4000~500 cm^−1^ wavenumber. (**B**) % transmittance change at 3200~3400 cm^−1^ following surface oxygen plasma treatment. Peak at ~1700 cm^−1^ corresponding to a C=C bond is observed in TMSPMA-modified PDMS surface. (**C**) Control: bare PDMS surface. Plasma: PDMS modified by oxygen plasma. TMSPMA: plasma-treated PDMS reacted with TMSPMA.

**Fig. S5.**
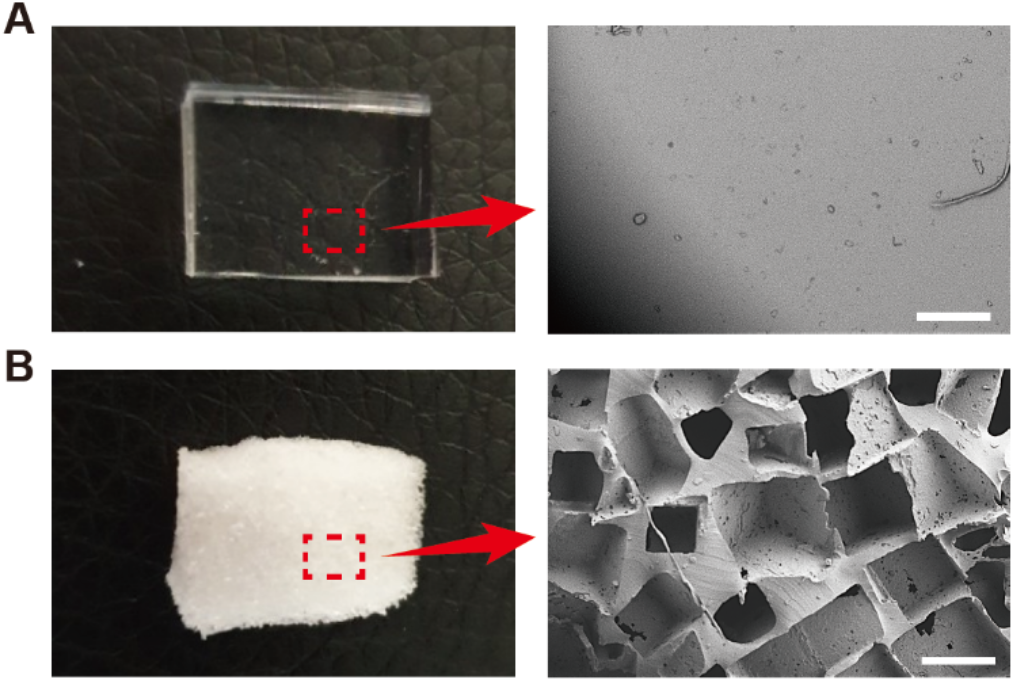
Scanning Electron Microscope images of PDMS microstructure. (**A**) Bare PDMS slab has a clear surface without any microstructure. (**B**) In contrast, microporous PDMS substrate shows interconnected salt crystal-shaped pores. Scale bar: 250 μm.

**Table S1.**
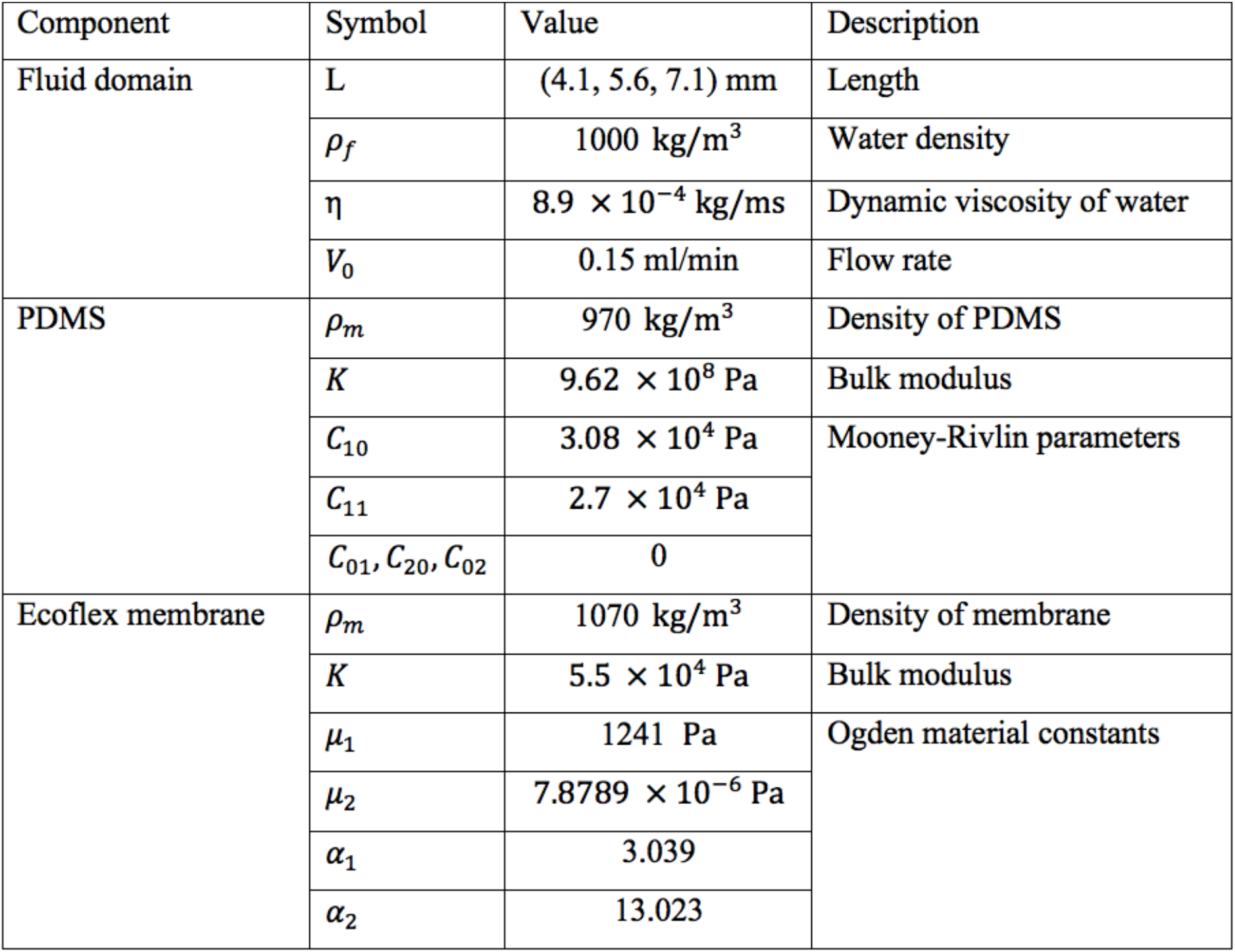
Physical and geometric parameters used in the numerical simulation.

## Supplementary Movies Caption

### Movie S1

Ecoflex 00-30 membrane inflates significantly, causing the entire PDMS structure to bend. The size of the rectangular channel in the balloon actuator is 7.1 × 1.8 × 0.15 mm (length × width × height).

### Movie S2

The maximum bending angle and the increase in rate of angle is proportional to channel length.

### Movie S3

Flapping/flexing motion of each hindwing is controlled separately by two syringes, and the independent flexing of each hindwing allows the DraBot to change traveling direction.

### Movie S4

Both hindwings flexed upward to unblock the air outlets of both forewings, leading to unidirectional airflow and forward propulsion. Maintaining the upward flap of the hindwings results in a straight motion.

### Movie S5

Left hindwing flexing generates net torque, resulting the bot to turn rightwards. A sustained right-turn signal leads to continuous clockwise motion.

### Movie S6

Net torque, generated by the right hindwing flexing, makes the bot turn leftwards. A sustained left turn signal leads to continuous counterclockwise motion.

### Movie S7

Long-term exploration of water surface by the bot through a combination of forward, left, and right turn motions.

### Movie S8

pH-sensitive self-healing of the A6ACA hydrogel facilitates welding of fore- and hindwing upon exposure to acid, which prevents the flapping of hindwing.

### Movie S9

Welding of fore and hindwings (left side) block the air outlet on left forewing, which results in DraBot making a continuous left turn, despite receiving an input signal for forward motion.

### Movie S10

When exposed to a temperature above 37 °C, the wing color changes from red-to-yellow, and reverts back when exposed to room temperature.

### Movie S11

The PDMS hydrophobicity along with the microstructure mediated increase in surface area allows microporous PDMS structure to imbibe hydrophobic oil while no such absorption is observed with water.

### Movie S12

The microporous PDMS structures present at the wings and abdomen absorb oil from the water when encountered. For better visualization, colorless PDMS was used to fabricate DraBot. The red color shows the absorption of oil.

